# Optimal bang–bang feedback for bursty gene expression

**DOI:** 10.1101/793638

**Authors:** Iryna Zabaikina, Pavol Bokes, Abhyudai Singh

## Abstract

Stochasticity in gene expression poses a critical challenge to the precise control of cellular function. In this paper we examine how precisely can a stochastically expressed protein attain a given target expression level. We consider a protein which is produced in bursts and which is able to control its expression via a negative feedback loop; we specifically focus on feedback of a bang–bang type which turns off the production of the protein whenever its concentration exceeds a given threshold. Using a piecewise deterministic mathematical formalism, we derive explicit expressions for the probabilistic distribution of the protein concentration, and for the mean square deviation from the target level. Employing a combination of analytic and numerical optimization, we identify the optimal value of the bang–bang threshold, in terms of minimising the deviation, and examine the dependence of the optimal value on the target level and the sub-threshold burst frequency. The systematic analysis allows us to formulate a number of quantitative and qualitative conclusions about the controllability of burst like gene expression. Finally, we outline directions for future research into the topic.

## I. INTRODUCTION

Bursty protein production is a major source of stochasticity (noise) in the expression of genetic information [1]. The bursts result from brief periods of intense transcriptional/translational activity, and are interspersed by long time spans of quiescence, during which the protein is degraded/diluted [2]. The amount of protein produced in a burst, referred to as the burst size, is a random quantity, as it depends on the length of the bursting period, which differs from one instance to another [3]. The duration of bursts is widely assumed to be memoryless [4]. Consequently, burst sizes are geometrically distributed [5] (if one uses a discrete copy number to measure the protein level) or exponentially distributed [6] if one uses a continuous concentration for the protein level. Here we will consider a continuous piece-wise deterministic framework with exponentially distributed burst sizes [7], [8], [9]. Importantly, this modelling framework explains large noise levels observed in gene expression at single cell level [10]. The question that we pose in this paper is one of optimality [11], [12], [13], [14]: we seek to determine how precisely we can target a prescribed protein concentration in the context of a highly random bursty protein expression.

Negative feedback is a control mechanism, which uses the information on a controlled quantity (here protein concentration) to counterbalance its deviations [15]. Feedback is ubiquitous in gene expression, with a specific class of protein, called transcription factors, being able to interact with the DNA to suppress their own expression [16]. Transcriptional feedback manifests itself through the regulation of burst frequency, meaning that high protein concentration leads to a decrease, and low concentration to an increase, in the frequency [17]. The dependence of the frequency on the protein concentration is given by the response function. Many studies use a Hill type response function [18]. Here we consider a step like function, which gives a positive value of frequency if the protein concentration is beneath a critical threshold, and a zero value if the value if the concentration exceeds the threshold. In line with the stated aim of the paper, we seek to determine the optimal value of the threshold which minimises the mean square deviation from the target level.

The outline of the paper is as follows. Section II formally introduces the model particulars. Section III provides explicit formulae for protein probability density functions, first in the unregulated case (Subsection III-A), and then in the feedback case (Subsection III-B); a special attention is given in Subsection III-C to the important case of infinite subthreshold burst frequency. Section IV deals with the problem of minimising the deviation from the target level, first for the unregulated framework, then for the infinite sub-threshold burst frequency (Subsection IV-A), and finally for finite values of the rate (Subsection IV-B). Section V concludes the paper with a discussion of its results.

## II. MODEL FORMULATION

The fundamental model for protein dynamics that we use throughout our work is a piecewise deterministic Markov process *x*(*t*) which decays deterministically (black portions of the trajectories in Fig. 1) but is produced stochastically in bursts (vertical red portions of the trajectories in Fig. 1). Decay is assumed to be simple exponential, i.e. the protein trajectory *x*(*t*) is proportional to e^−*γt*^ inside any time interval which does not contain a burst event. In the absence of feedback, bursts of production occur randomly in time with frequency (or rate) *a* per unit time; the inclusion of feedback will be dealt with below. Whenever a burst event occurs, the protein concentration is discontinuously increased by the burst size, which is randomly drawn from the exponential distribution with mean *b*. By measuring time in units of protein lifetime, we can achieve, without any loss of generality, that *γ* = 1. The parameter *a* then assumes the meaning of normalized burst frequency, and gives the expected number of bursts occurring per typical protein lifetime.

**Fig. 1:**
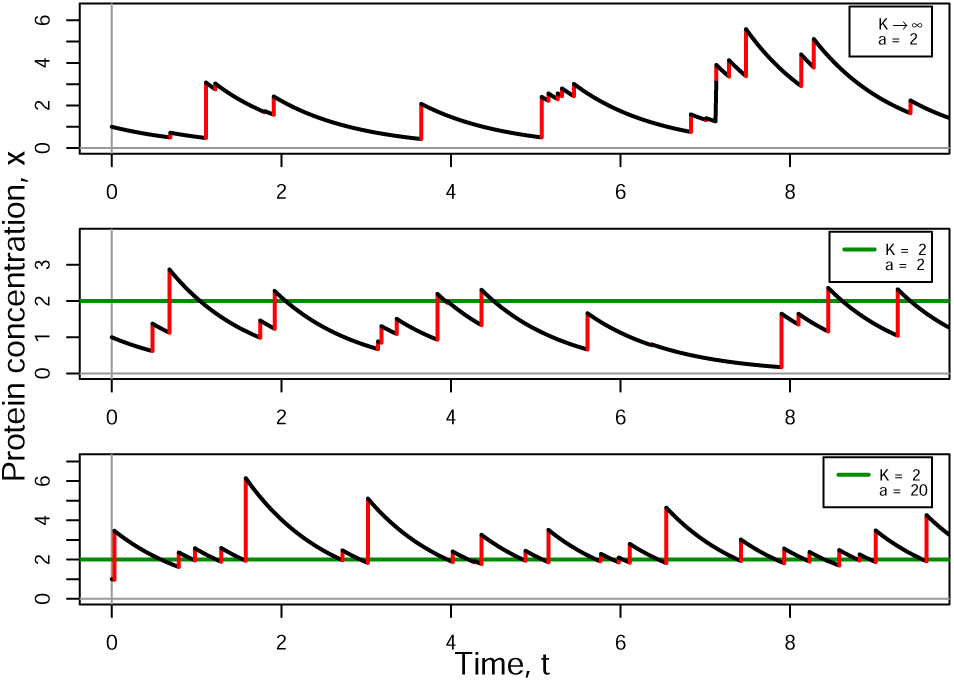
Sample trajectories of gene expression without any regulation (the upper panel) and subject to negative feedback with the *K* = 2 (green horizontal line). The burst size is fixed to *b* = 1 and burst frequency is *a* = 2 (upper and middle panels) and *a* = 20 (bottom panel).

Extending our model by feedback in burst frequency, we allow the production rate to depend on the current protein concentration so that

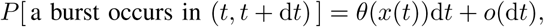

where *θ*(*x*) is a response function which determines the production rate in the presence of feedback. We will focus on negative feedback only, meaning that we consider non-increasing functions *θ*(*x*). Specifically, the Hill function [16]

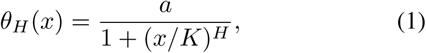

where *a, H*, and *K* are parameters explained below, has been widely used in literature and justified in terms of cooperative binding of a protein molecule to its gene’s promoter. The parameter *K* gives the concentration required to reach half the maximum burst frequency; it corresponds to the dissociation constant of the protein–promoter binding [19]. Interestingly, by taking *K* to infinity one recovers the unregulated model as introduced earlier in this section. The parameter *H*, which is referred to as the Hill coefficient, indicates how steeply feedback reacts to changes in protein concentration (Fig. 2), and directly corresponds to the co-operativity in the underlying promoter–protein independent interaction. The parameter *a* here takes the role of maximal burst frequency, which is achieved in the absence of the self-repressing protein.

**Fig. 2:**
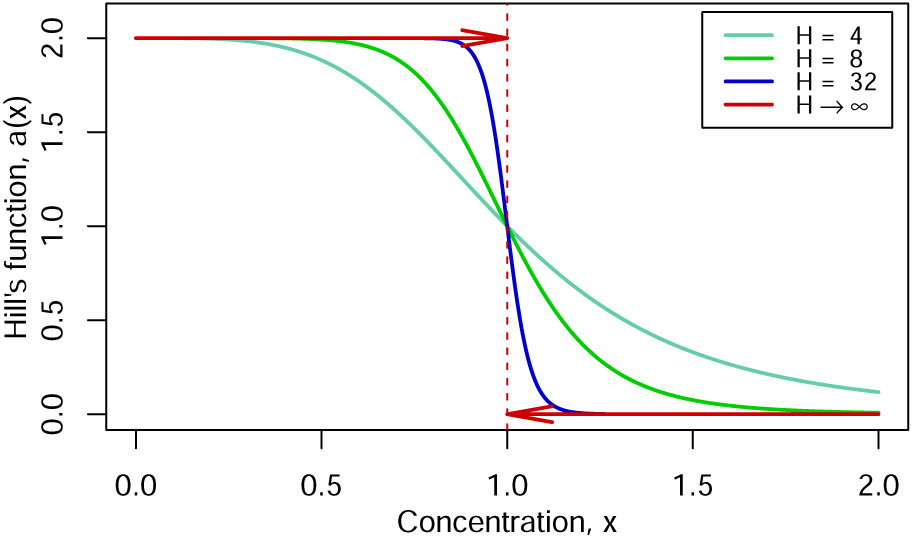
Examples of Hill functions (1) with different choices of Hill coefficients *H*. As *H* increases to infinity, the function develops at *x* = *K* a jump discontinuity. The other parameters of (1) are set to *a* = 2 and *K* = 1.

In the limit of very large Hill coefficients, we find that (cf. Fig. 2)

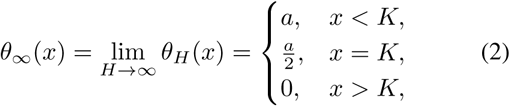

which we hereafter refer to as bang–bang feedback. In case of bang–bang type feedback, the protein exponentially degrades, and bursts cannot occur, as long as the concentration level is higher than *K*; once *x* falls beneath *K*, bursts occur with a constant frequency *a*.

## III. Protein concentration distribution

The (stationary) distribution of protein concentration is obtained as a steady-state solution of a master equation associated to the process described in the previous section. The master equation here takes the form of a probability conservation law [20]

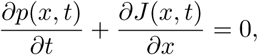

in which the probability flux is shaped by the model’s particulars and here given by [9]

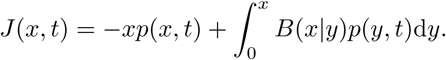

In the probability flux *J* (*x, t*), the first term corresponds to the decay of protein level due to natural protein degradation, and the second term corresponds to an instant growth of the protein level in bursts; *B*(*x*|*y*) is the burst kernel, which defines the conditional probability that concentration will jump from a given protein level *y* to beyond *x* after burst.

### A. Unregulated gene expression

The burst kernel in case of unregulated gene expression is given by the product

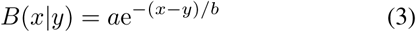

of burst frequency *a* and the probability e^−(*x*−*y*)*/b*^ that the burst size is larger than the difference *x* − *y*.

The steady-state of protein concentration is reached when system dynamics do not change with time, so that *∂p*(*x, t*)*/∂t* = 0, from which it follows that the probability flux *J* (*x, t*) must be constant with respect to *x*; and since we do not admit a nonzero flux of probability mass from infinity, this constant has to be equal to zero. Thus the equation of stationary probability density function satisfies *J* = 0, i.e.

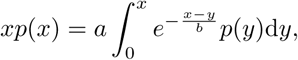

which is an Volterra integral equation of the second kind with a difference kernel, for which one can obtain *p*(*x*) using Laplace transformation approach [21]. This approach involves transforming the integral equation into a differential equation for the Laplace image, and subsequently returning to the original space.

Performing this solution approach one finds that [7]

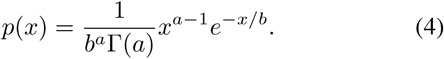

It means that in the unregulated gene expression the protein concentration *x* ∼ Γ(*a, b*), i.e. it is gamma-distributed with shape *a* and scale *b*. From this follows, in particular,

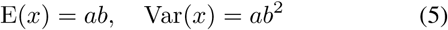

for the protein mean and variance.

### B. Regulated gene expression

The negative regulation kernel is given by the product of the response function *θ*(*y*) and the probability of a burst exceeding the size of *x* − *y*, i.e.

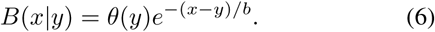

Note that while this kernel is no longer a difference kernel, it is still a product kernel, and, as such, the associate integral equation

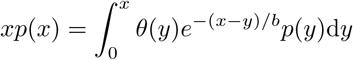

admits an explicit solution [22]. In order to find it, we pull out from under the integral sign the exponential *e*^−*x/b*^, differentiate the equation with respect to *x* and apply the Leibnitz integral rule; this yields an ODE

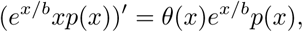

the general solution of which is

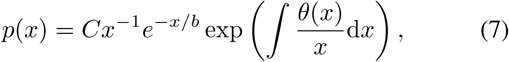

in which *C* is an integration constant. The primitive function in the argument of the exponential can easily be solved for the piecewise constant *θ*(*x*) of the bang–bang regulation (2). Since the bang–bang response function features a discontinuity at *x* = *K*, the primitive function in the exponential, as well as the probability density function (PDF) *p*(*x*) itself, are nonsmooth at *x* = *K* (Fig. 3); to the left and to the right of the point of smoothlessness the density is given by separate expressions

**Fig. 3:**
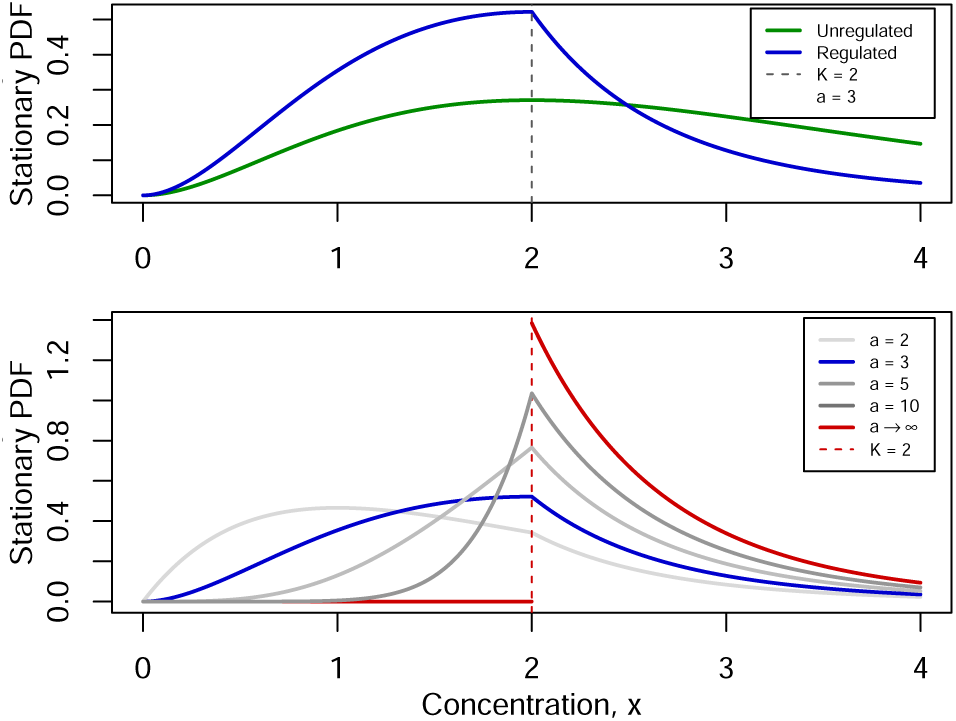
*Upper:* The PDF of protein concentration governed by negative feedback (blue line) in comparison with the PDF of unregulated gene expression (green line). *Lower:* PDFs with finite sub-threshold burst frequencies (grey and blue lines) and with the infinitely high value (red line) are shown; the probability of *x* being less than *K* decreases as *a* increases. We set *b* = 1 throughout.

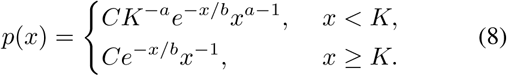

The condition that the distribution density must be normalized to one fixes the value of the normalisation constant to

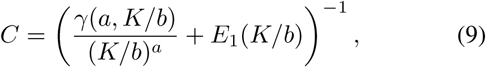

where *γ*(*a, z*) is the lower incomplete gamma function [23] is defined by

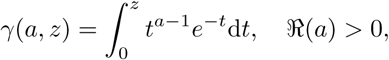

and *E*_1_(*z*) is the exponential integral defined on the complex plane by

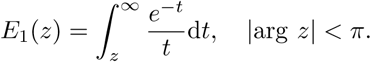

Integrating the density multiplied by the factor *x*, we obtain

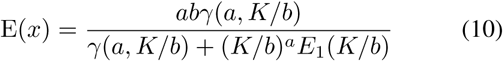

for the expected protein concentration at steady state.

### C. Infinitely high frequency case

Let us investigate in greater detail the limiting case of burst frequency *a* that is infinitely high; in such case it is expected that, if the protein level falls below *K*, a burst occurs almost immediately; therefore the concentration is higher than K almost surely. Since only a single parameter has been taken to a limit but the fundamentals of the model remain unaltered, the stationary probability density function for protein concentration can be derived by taking a limit as *a* → *∞* of (8).

Provided that the protein concentration *x* is less than the dissociation constant *K*, we have

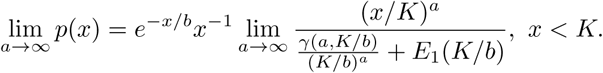

The same approach for the other case, occurring when the protein concentration *x* is above *K*, yields

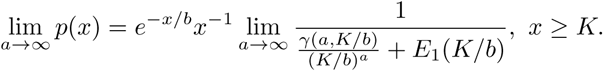

The lower incomplete gamma function has series expansion [23]

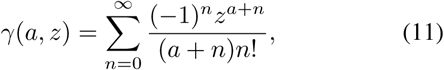

from which

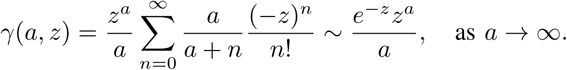

In particular it follows that

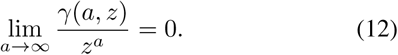

The limiting stationary density function therefore given by

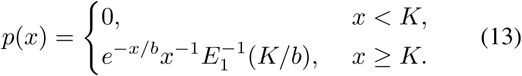

The change in the form of the protein probability density function in response to increasing the sub-threshold frequency is illustrated on lower panel of Fig. 3, with the limiting case (13) shown in red. The mean protein concentration is 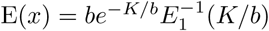

## IV. Optimization problem

Here we use the theoretical results of the preceding sections to study the mean squared difference

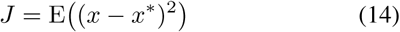

between the steady state protein level and a prescribed target level *x*^*^. The mean square difference can be written as

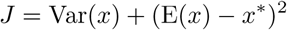

in terms of the protein mean and variance.

In case of unregulated gene expression, the mean and variance are given by (5), implying that

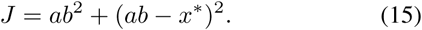

The nonnegative value *a* = *a*_opt_ that minimises the above expression and the minimum *J*_opt_ = *J* (*a*_opt_) are given by

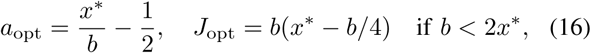

or

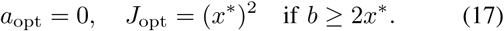

We will see that, by employing the bang–bang feedback, much lower values can be achieved.

In the presence of bang–bang feedback, we write the mean square difference as

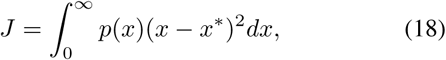

in which the probability density function is given by (8) if *a* < *∞* or (13) if *a* = *∞*. The value of *a* is thereby assumed to be a fixed constant. We shall consider the mean squared difference (18) to be a function of the dissociation constant, i.e. *J* = *J* (*K*), seeking to determine the value *K*_opt_ at which the cost function *J* achieves an optimal (minimal) value *J*_opt_ = *J* (*K*_opt_).

### A. Infinitely high frequency case

Substituting into (18) the functional form of the PDF *p*(*x*) in the infinitely high sub-threshold burst frequency (13), we arrive at

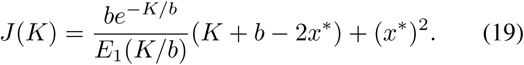

The graph of this function, including the location of its minimum, is shown in the Fig. 4. The graph includes the behaviour of *J* (*K*) only for right-sided neighbourhood of *K* = 0 because both the definition of exponential integral and the interpretation of the model do not admit negative values of the dissociation constant.

**Fig. 4:**
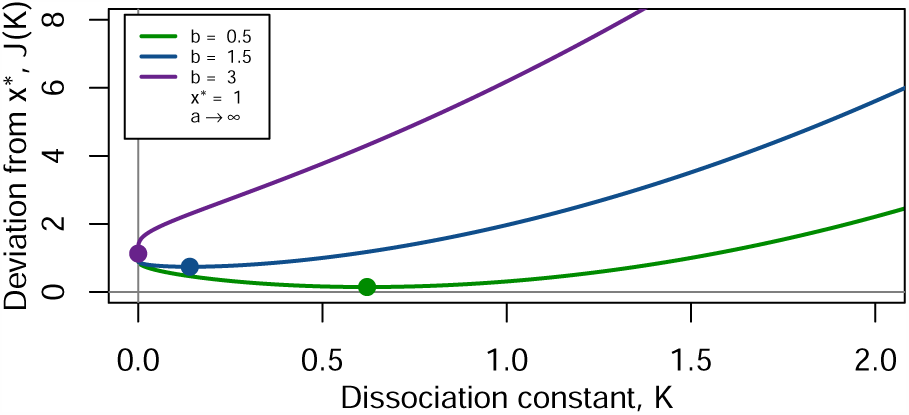
The cost function *J* (*K*) for *a* = *∞* and a selection *b* values. The markers indicate the coordinates of the minimizer *K*_opt_ and the minimum *J*_opt_.

Differentiating the cost function (19) yields

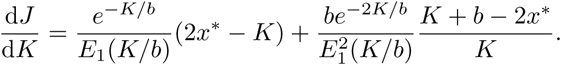

For low values of the threshold *K* we have an asymptotic approximation

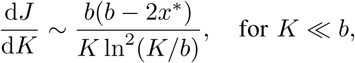

which implies, in particular, that

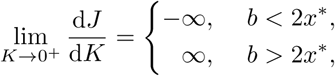

so that the cost function is locally decreasing for 0 < *K* ≪*b* if *b* < 2*x*^*^ and locally increasing if *b* > 2*x*^*^ (cf. Fig. 4). The local behaviour for small positive values of *K* has important implications for the global minimum of the function *J* (*K*): if *b* > 2*x*^*^, the local minimum is achieved at *K*_opt_ = 0, whereas for *b* < 2*x*^*^, the local minimum *K*_opt_ is found within the interval 0 < *K* < *∞* as the solution of the equation

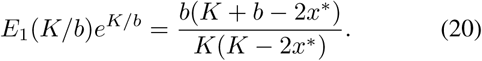

The dependence of *K*_opt_ on the mean burst size *b* is shown for *a* = *∞* in magenta colour in the right panel of Fig. 5b; note that *K*_opt_ = 0 for *b* > 2*x*^*^ and that *K*_opt_ ∼ *x*^*^ − *b* for *b* ≪*x*^*^. The left panel of Fig. 5b shows the minimal mean square deviation *J* (*K*_opt_) as function of the burst size *b*. If *b* > 2*x*^*^, then *K*_opt_ = 0 implies *x ≡* 0, so that *J* (*K*_opt_) = E(*x* − *x*^*^)^2^ = (*x*^*^)^2^. As the burst size *b* decreases to zero, the minimal mean square deviation becomes progressively smaller with asymptotics *J* (*K*_opt_) ∼ *b*^2^, *b* ≪*x*^*^ (Fig. 5b, left, magenta colour). Note that this quadratic decrease is faster than the linear one exhibited by the minimal deviation in the unregulated case (16).

**Fig. 5:**
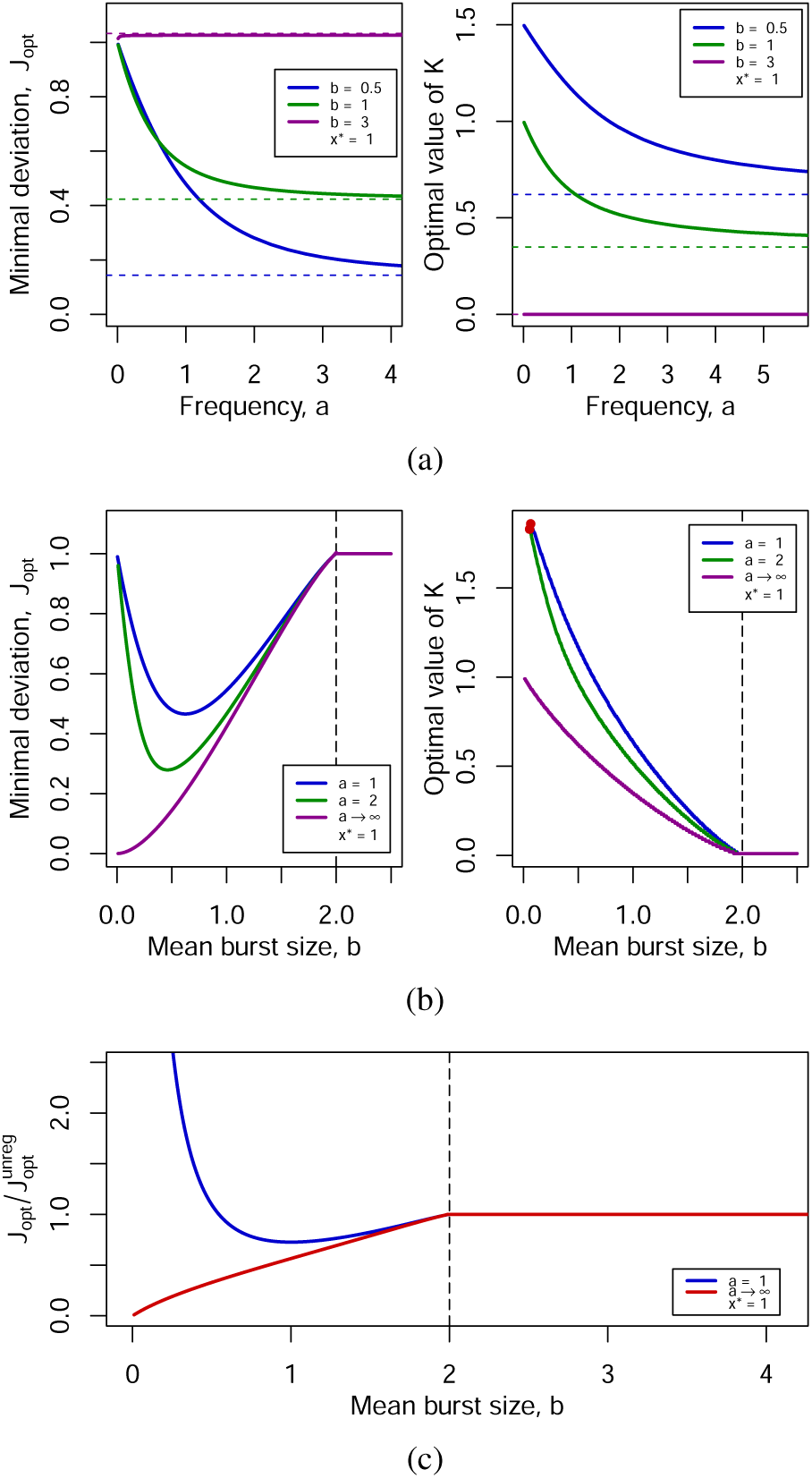
(a) The minimal square deviation *J*_opt_ (*left*) and the minimizer *K*_opt_ (*right*) as functions of *a* for selected *b*. The horizontal dashed lines indicate the limit values as *a* → *∞*. In the opposite limit *a* → 0 (protein production is turned off) we find *K*_opt_ → (2*x*\* − *b*)_+_ and *J*_opt_ → (*x*^*^)^2^. (b) The minimal square deviation *J*_opt_ (*left*) and the minimizer *K*_opt_ (*right*) as functions of *b* for selected *a*. The vertical line refers to the critical value *b* = 2*x*\*; red dots indicate the values of *b* below which the calculation of *K*_opt_ is precluded by numerical indeterminacy issues. (c) The optimal cost of the regulated feedback *J* (*K*_opt_) normalized by that of the unregulated one *J* (*a*_opt_) (16)–(17).

The left-hand side of (20) with the special function *E*_1_(*K*) carries with it an additional difficulty when solving this equation; it will be sufficient for us to solve the equation numerically with high precision. To do this, we use the package `pracma` of the language R for statistical computing [24], which provides all the necessary functionality. There are two ways to find solution *K*_opt_: the first is to find directly the minimal value of the cost function; the second is to find the point, at which the difference between the left and right hand sides of (20) is the smallest. In practice, the first method appears to be more robust and accurate.

### B. Finite frequency case

Here we consider a more general case of the gene expression regulation, in which *a* is any finite positive number. Inserting the functional form (8) of the PDF *p*(*x*) into the cost function (18), we find that the mean square deviation of the protein concentration *x* from *x*^*^ is given by a sum of two terms

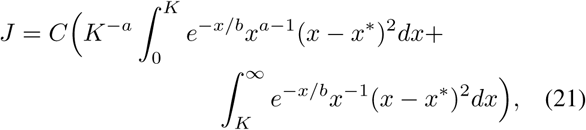

where *C* is the normalisation constant (9). After applying certain transformations and expressing the integrals in terms of special functions, we arrive at a representation

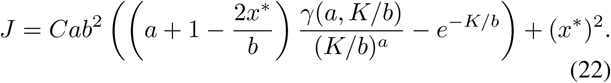

Aiming to find the minimum of the cost function, we differentiate (22) with respect to *K*, obtaining

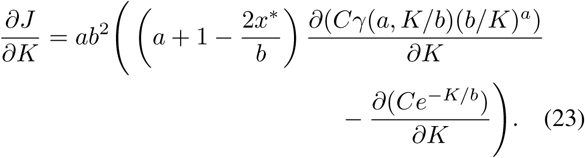

The sought-after optimal value of *K*_opt_, at which the protein concentration *x* minimally fluctuates around the required level *x*^*^, satisfies the equation

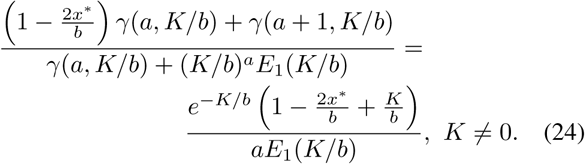

In Fig. 5, we exemplify the behaviour of the cost function *J* (*K*) and study the dependence of the optimiser *K*_opt_ on the upregulated burst rate *a* and the burst size *b*. The information conveyed by the figure is discussed in detail in the next section.

## V. Discussion

In this paper we considered the question of finding optimal parameters in bursting gene expression models so as to minimise the cost function *J* (14) which is defined as the square distance from a prescribed protein level *x*^*^. First, we examined this question in the context of unregulated gene expression occurring with burst frequency *a* and mean burst size *b*. The steady state distribution is the gamma distribution with shape *a* and scale *b*; the mean square difference from the target is given by a simple multinomial of the model parameters (15). In our analysis we fixed *b* and *x*^*^ and sought to find an optimal *a* = *a*_opt_. Elementary analysis showed that a positive optimal burst frequency *a*_opt_ which minimises the deviation exists only if the mean burst size is less than twice the target level (*b* < 2*x*^*^); otherwise it is optimal to shut off the production of protein by setting the optimal burst frequency *a*_opt_ to zero. Thus, the elementary analysis of the unregulated case demonstrates that large burst sizes limit the ability of a gene to achieve relatively low targets.

After examining the unregulated case, we turned our attention to the case of bang–bang feedback. Its performance is strongly dependent on the choice of the critical threshold (also referred to as the dissociation constant) *K* at which the gene expression gets shut down. For a fixed target level *x*^*^, sub-threshold burst frequency *a* (which can be finite or infinite) and mean burst size *b*, we calculated the value *K* = *K*_opt_ which minimises the mean square deviation from *x*^*^. We found that, similarly as in the unregulated case, a positive optimal threshold is available only if the mean burst size is smaller than twice the target level (*b* < 2*x*^*^); otherwise it is optimal to shut off gene expression completely by setting the threshold *K*_opt_ to zero (Fig. 5b). Decreasing the sub-threshold burst frequency *a* leads to an increase in both the optimal threshold *K*_opt_ and the optimal deviation *J*_opt_ = *J* (*K*_opt_) (Fig. 5a). If the burst size is very small, then a protein with a finite value of burst freqency *a* will fail to reach the target *x*^*^ whatever the value of the feedback threshold *K* may be; this explains both the large values of *J*_opt_ as well as indeterminacy of *K*_opt_ for *a* < *∞* and *b* ≪*x*^*^ (Fig. 5b). In light of these observations, the infinite sub-threshold burst frequency (*a* = *∞*) is deemed to be best performing among bang–bang feedbacks. Additionally, a protein with an optimal bang–bang feedback with *a* = *∞* outperforms a protein without regulation in terms of reaching the target *x*^*^ (Fig. 5c).

In addition to providing a number of observations for the optimisation problem, our analysis contributes to previous results by establishing in Equations (8) and (13) the form of the steady state probability density function for a protein subject to bang–bang burst frequency feedback. In the future steps of our work, we intend to use optimal-control techniques, such as the Pontryagin maximum principle, to establish the optimality of bang–bang feedback among a wider class of response functions. Additionally, we intend to extend our analysis to other types of gene-expression feedback, such as feedback in burst size [25] and protein stability [26].

## Notes

PB is supported by the Slovak Research and Development Agency under the contract No. APVV-18-0308, by the VEGA grant 1/0347/18, and the EraCoSysMed project 4D-Healing. AS is supported by the National Science Foundation grant ECCS-1711548.

